# Engineered Migrasomes: Harnessing Core Migrasome Machinery and Hypotonic Shock to Develop a Robust and Thermally Stable Vaccine Platform

**DOI:** 10.1101/2024.03.13.584850

**Authors:** Dongju Wang, Haifang Wang, Wei Wan, Zihui Zhu, Takami Sho, Yi Zheng, Xing Zhang, Longyu Dou, Qiang Ding, Li Yu, Zhihua Liu

## Abstract

The increasing ability of pathogens and tumor cells to evade immune detection underscores the urgent need for novel vaccine platforms leveraging diverse biological mechanisms. Additionally, logistical challenges associated with cold-chain transportation significantly limit vaccine accessibility, especially in resource-limited regions. Recently, we identified migrasomes, specialized organelles generated during cell migration, which are inherently stable and enriched with immune-modulating molecules. To address the low yield of natural migrasomes, we engineered migrasome-like vesicles (eMigrasomes) using hypotonic shock combined with cytoskeletal disruption to enhance vesicle formation. The biogenesis of eMigrasomes relies on the core migrasome machinery, faithfully recapitulating the biophysical attributes of native migrasomes while significantly improving production efficiency. We demonstrate that eMigrasomes loaded with a model antigen elicit potent antibody responses and maintain structural integrity and immunogenic potential at room temperature. Furthermore, eMigrasomes displaying the SARS-CoV-2 Spike protein induce robust humoral immune responses, providing effective protection against viral infection. Our findings highlight the potential of utilizing migrasome biology and hypotonic shock-driven vesicle generation as an innovative, stable, and broadly accessible vaccine platform.

## Introduction

To face the imminent challenges of emerging infectious diseases, and to realize the not-yet-fulfilled promise of cancer vaccines, new types of vaccination platforms based on different basic biological principles are urgently needed. The huge success of mRNA vaccines in the fight against the Covid-19 pandemic further supports the importance of developing vaccine platforms based on different underlying biological mechanisms ^1–4^. However, the application of mRNA-based vaccines, or other forms of traditional vaccine, has been hampered by the requirement for complicated cold-chain transportation, in which the materials are constantly maintained at low temperature ^5, 6^. For most parts of the world, especially the global south, it is vitally important to develop vaccines which do not require cold-chain transportation.

Migrasomes are recently discovered organelles of migrating cells ^7^. During migrasome formation, long membrane tethers named retraction fibers are pulled out at the trailing edge of migrating cells by the force generated by the movement of the cells. Migrasomes are formed on retraction fibers by a complicated process involving multiple components ^8–10^. In the final expansion step, the growth of migrasomes is driven by assembly of nanometer-scaled tetraspanin-enriched microdomains (TEMs) into micrometer-scaled tetraspanin-enriched macrodomains (TEMAs) ^9, 11^. Thus, migrasomes are highly enriched with components of TEMs — such as tetraspanins, integrins, and cholesterol — and formation of migrasomes is dependent on the presence of these molecules. For example, overexpression of Tspan4 can promote migrasome formation ^9^, while removing cholesterol can block migrasome formation. Theoretical modelling, *in vitro* reconstitution of migrasome formation, and membrane stiffness measurement by atomic force microscopy have revealed mechanistic insights into migrasome formation. These approaches showed that tetraspanin- and cholesterol-enriched macrodomains have highly elevated membrane stiffness, which is the key factor to drive the bulging of retraction fibers into migrasomes; as a result, migrasomes are highly rigid ^9^. More recently, it was shown that assembly of TEMs can repair damaged membranes by restricting the spread of membrane rupture. In liposomes containing Tspan4 and cholesterol, detergent-induced membrane damage can be rapidly repaired, and Tspan4-embedded liposomes are highly resistant to damage ^12^.

One defining feature of tetraspanin-enriched microdomains is the enrichment of immune-modulating molecules such as members of the immunoglobulin superfamily (IgSF) ^13–15^. This group of molecules contains many key signaling molecular complexes which regulate the immune response, including antigen receptors, antigen-presenting molecules, co-receptors, antigen receptor accessory molecules, and co-stimulatory or inhibitory molecules ^16^. Since migrasomes are largely composed of TEMs, members of the IgSF are also enriched in migrasomes. This property, in addition to the high stability of migrasomes, prompted us to explore the possibility of developing a migrasome-based vaccine.

One key obstacle to developing a migrasome-based delivery system and migrasome-based vaccines is the very low yield of migrasomes. To generate migrasomes, cells must be grown at very low density to allow them to migrate, and a migrating cell can only generate a relatively low number of migrasomes in a process that takes hours to finish ^7^. In this study, by using the biophysical insights we gained from investigating migrasome formation, we successfully overcome this key obstacle. We developed a method which mimics the key biophysical features of migrasome formation while vastly improving the yield of migrasome-like vesicles. We named these vesicles as engineered migrasomes (eMigrasomes). Using eMigrasomes loaded with the model antigen ovalbumin (OVA), we successfully demonstrate that eMigrasomes are a highly effective, room temperature-stable vaccine platform which generate a strong immunoglobulin G (IgG) response. Finally, we demonstrate that eMigrasomes loaded with SARS-CoV-2 Spike protein (S protein) can generate a strong protective humoral response against SARS-CoV-2. In summary, our study demonstrates that the eMigrasome-based vaccine platform is stable at room temperature, highly effective, and uses an unconventional immunological mechanism, which could be useful in certain scenarios where conventional vaccine platforms are less successful.

## Results

By serendipity, we found that hypotonic shock induces rapid formation of migrasome-like structures on retraction fibers in Tspan4-GFP-expressing cells (Fig 1a). Similar phenomena had also been observed in recent studies^17, 18^. 10 seconds after hypotonic shock, the Tspan4 signal started to become enriched on retraction fibers. These Tspan4-enriched domains then grew into migrasome-like structures. After reaching its peak intensity, the Tspan4-GFP signal started to diffuse away from the migrasome-like structure, which was accompanied by shrinkage of the migrasome-like structure. 440 seconds after hypotonic shock, most of the hypotonic shock-induced migrasome-like structures disappeared (Fig 1a, Supplementary Video 1). We found that the size of the migrasome-like structures negatively correlated with osmolarity: the lower the osmolarity, the larger the migrasome-like structures (Fig 1b, 1c). To study the role of osmolarity in the formation of migrasome-like structures, we set up an imaging protocol which allowed us to carry out live-cell imaging while lowering the osmolarity in a step-wise manner (Fig 1d). We found that step-wise application of the hypotonic shock significantly increased the duration time of the migrasome-like structures. In cells undergoing 5 steps of hypotonic shock, 50% of the migrasome-like structures were still stable 6 minutes after the final step of hypotonic shock; in contrast, in cells undergoing one step of hypotonic shock, 50% of the migrasome-like structures shrunk back or fused with their neighbors within 1.5 minutes. (Fig 1e, 1f, Supplementary Video 2, 3).

**Figure 1.**
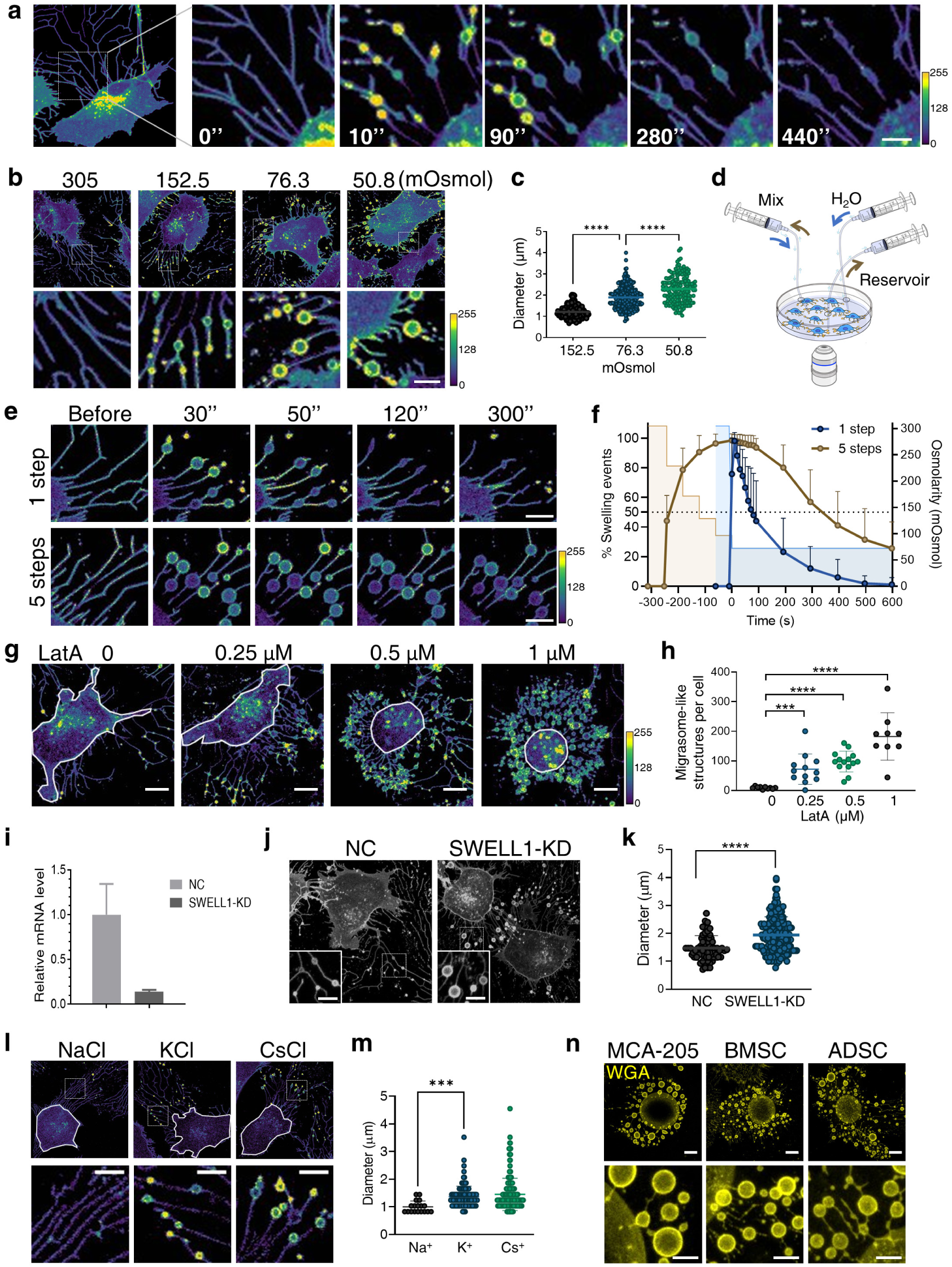
Hypotonic stimulation induced migrasome-like structures. (a-m) The biogenesis of migrasome-like structures in NRK cells stably expressing Tspan4-GFP. In these images, the fluorescence intensity of Tspan4-GFP is shown in a color map scale from purple (low) to yellow (high). (a) Image series of the biogenesis of migrasome-like structures. Cells were treated with hypotonic Dulbecco’s phosphate-buffered saline (DPBS) with an osmolarity of 76.3 mOsmol and imaged using a confocal microscope. Scale bar, 10 μm. (b) Representative confocal images of cells treated with DPBS with various osmolarities. 305 mOsmol represents an isotonic condition in which no migrasome-like structures were observed. DPBS diluted to 152.5, 76.3 or 50.8 mOsmol was used to achieve hypotonic stimulations of different magnitude. The size of migrasome-like structures increased as the osmolarity was reduced. Scale bar, 5 μm. (c) For each migrasome-like structure in (b), the whole lifetime, including the growth and shrinkage, was recorded by time-lapse imaging. The largest diameter reached during the lifetime was measured. The average diameter of migrasome-like structures increased significantly as the osmolarity was reduced. For hypotonic stimulation at 152.5, 76.3 or 50.8 mOsmol, n = 194, 261, 165 migrasome-like structures, respectively. (d) Illustration of the experimental setup for real-time imaging of the induction of migrasome-like structures. (e) Image series of cells treated with hypotonic DPBS in two different approaches. The osmolarity of DPBS was reduced to 76.3 mOsmol by either one single step (upper panel) or five steps with 1 min intervals (lower panel). For the step-wise reduction, the osmolarity was lowered by 25% in each step. Imaging time-points were counted from the final stimulation step. Migrasome-like structures induced by the stepwise protocol showed significantly enhanced stability. Scale bar, 5 μm. (f) Statistical analysis of growth curves of migrasome-like structures in (e). For single step stimulation, n = 16 cells; for step-wise stimulation, n = 13 cells. (g) Representative confocal images showing the effect of Latrunculin A (LatA) treatment on the biogenesis of migrasome-like structures. Cells were pre-incubated with 0, 0.25, 0.5 or 1 μM LatA for 10 mins and then treated with a five-step hypotonic stimulation as described in (e). LatA enhanced the biogenesis of migrasome-like structures in a dose-dependent manner. The white line indicates the boundary of the cell body. Scale bar, 10 μm. (h) Statistical analysis of the number of migrasome-like structures per cell in (g). n = 9-14 cells. (i) Relative mRNA level of SWELL1 analyzed by qPCR. SWELL1 expression was significantly reduced in SWELL1-knockdown (KD) cells compared to control cells. (j) Representative confocal images of control or SWELL1-KD cells. Cells were treated with a five-step hypotonic stimulation at 2 min intervals. The osmolarity was reduced by 1/6 in each step. Migrasome-like structures are shown in the inserts. Scale bar, 5 μm. (k) Statistical analysis of the diameter of migrasome-like structures in (j); n=75 for control cells and 104 for SWELL1-KD cells. (l) Representative confocal images showing the effect of extracellular cations on the biogenesis of migrasome-like structures. Before stimulation, culture medium was replaced by modified DPBS in which either Na^+^, K^+^ or Cs^+^ was the only cation source. Cells were then treated with a five-step hypotonic stimulation at 2 min intervals. The osmolarity was reduced by 1/6 in each step. Scale bar, 5 μm. (m) Statistical analysis of the diameter of migrasome-like structures in (l); n = 18 for Na^+^, 119 for K^+^ and 203 for Cs^+^. (n) Multiple primary cell types and cell lines are capable of producing eMigrasomes. Cells were stained with WGA-AF488 (Thermo, W11261) after hypotonic induction of eMigrasome with our protocol. Z-stack image series were captured and sum-slices projections were applied. Scale bar, 10 μm (upper panels) and 5 μm (lower panels). For all statistical analyses in this figure, *P* values were calculated using a two-tailed unpaired nonparametric test (Mann-Whitney test). *P* value<0.05 was considered statistically significant. *** *P*<0.001. **** *P*<0.0001.

We realized that the amount of migrasome-like structures generated by hypotonic shock depends on the number of retraction fibers. We also realized that the majority of the plasma membrane can be the source of membrane for migrasome-like structures. We reasoned that if we shrink the cells by disrupting the cytoskeleton, the shrinkage will cause the retraction of the cell edge toward the center. At the same time, the cells will be adhering to the bottom of the culture plate at various points by focal adhesion. These adhesion sites will keep the plasma membrane in place, thus serving as anchor points for generation of membrane tethers. If this scenario is true, contraction of the cell will generate large numbers of membrane tubes in a way similar to retraction fiber formation during migration. To test this hypothesis, we treated cells with different doses of latrunculin A (LatA), a reagent widely used for disruption of microfilaments, before applying the step-wise hypotonic shock. As expected, we found that treating cells with LatA caused shrinking of the cells. We also observed the massive formation of membrane tethers in the area which was occupied by the cell before shrinkage. Importantly, we observed significantly enhanced formation of migrasome-like structures from these newly formed membrane tethers in a LatA dose-dependent manner (Fig 1g, 1h).

It is well established that cells can counter a change of osmolarity with regulated volume change ^19^. To test whether regulated volume change can affect the formation of migrasome-like structures, we knocked down SWELL1, a key component of the volume-regulated anion channel which maintains a constant cell volume in response to osmotic changes ^20, 21^. We found that knockdown of SWELL1 significantly enhanced the formation of migrasome-like structures. This result suggests that formation of migrasome-like structures can be enhanced by reducing the capacity of cells to regulate their volume during osmolarity change (Fig 1i - 1k).

Under normal physiological conditions, the extracellular space typically contains a substantial amount of sodium. Throughout all the aforementioned experiments, the buffer employed predominantly consisted of sodium as the prevailing cation. Cations are known for their role in regulation of cell volume. Next, we tested the effect of different cations on the formation of migrasome-like structures. We reasoned that if different cations have different abilities to regulate cell volume during osmolarity change, we may be able to find an easy way to attenuate the regulated cell volume change, thus promoting the formation of migrasome-like structures. To do that, we first replaced the medium with isotonic buffers containing different cations, then we added water step-wise to reduce the osmolarity. We found that indeed different cations have different abilities to promote the generation of migrasome-like structures. Substitution of sodium with equal molar potassium, cesium or choline significantly enhanced the generation of migrasome-like structures (Fig 1l, 1m).

Tspan4 is the key protein that promotes migrasomes formation, and all the experiments described above were carried out in Tspan4-GFP-expressing cells. To test whether Tspan4 can promote the formation of migrasome-like structures, we treated mCherry- or Tspan4-mCherry-expressing cells with step-wise hypotonic shock, and then observed the formation of migrasome-like structures by WGA staining, which labels migrasomes effectively ^22^. We found that indeed Tspan4-mCherry significantly enhanced the formation of migrasome-like structure (Fig 2a, 2b). Similar results held for Tspan1 and CD82, which are also migrasome-promoting tetraspanins (Fig 2c, 2d, S1a, S1b). Previously, we reported that cholesterol is essential for migrasome formation. We treated cells with methyl-beta-cyclodextrin (MβCD), which selectively extracts cholesterol from the plasma membrane. We found that addition of MβCD rapidly destroyed the readily formed migrasome-like structures (Fig 2e, 2f). This suggests that, similar to migrasomes, the structural integrity of migrasome-like structures depends on cholesterol. Recently, we reported that formation of migrasomes is dependent on SMS2. We found that treating cells with SMS2-IN-1, a selective inhibitor of SMS2, significantly inhibits the formation of migrasome-like structures (Fig 2g, 2h). Finally, we tested whether the formation of migrasome-like structures can occur in cells other than NRK cells. Indeed, all the cells we tested were able to support the formation of migrasome-like structures (Fig 1n).

**Figure 2.**
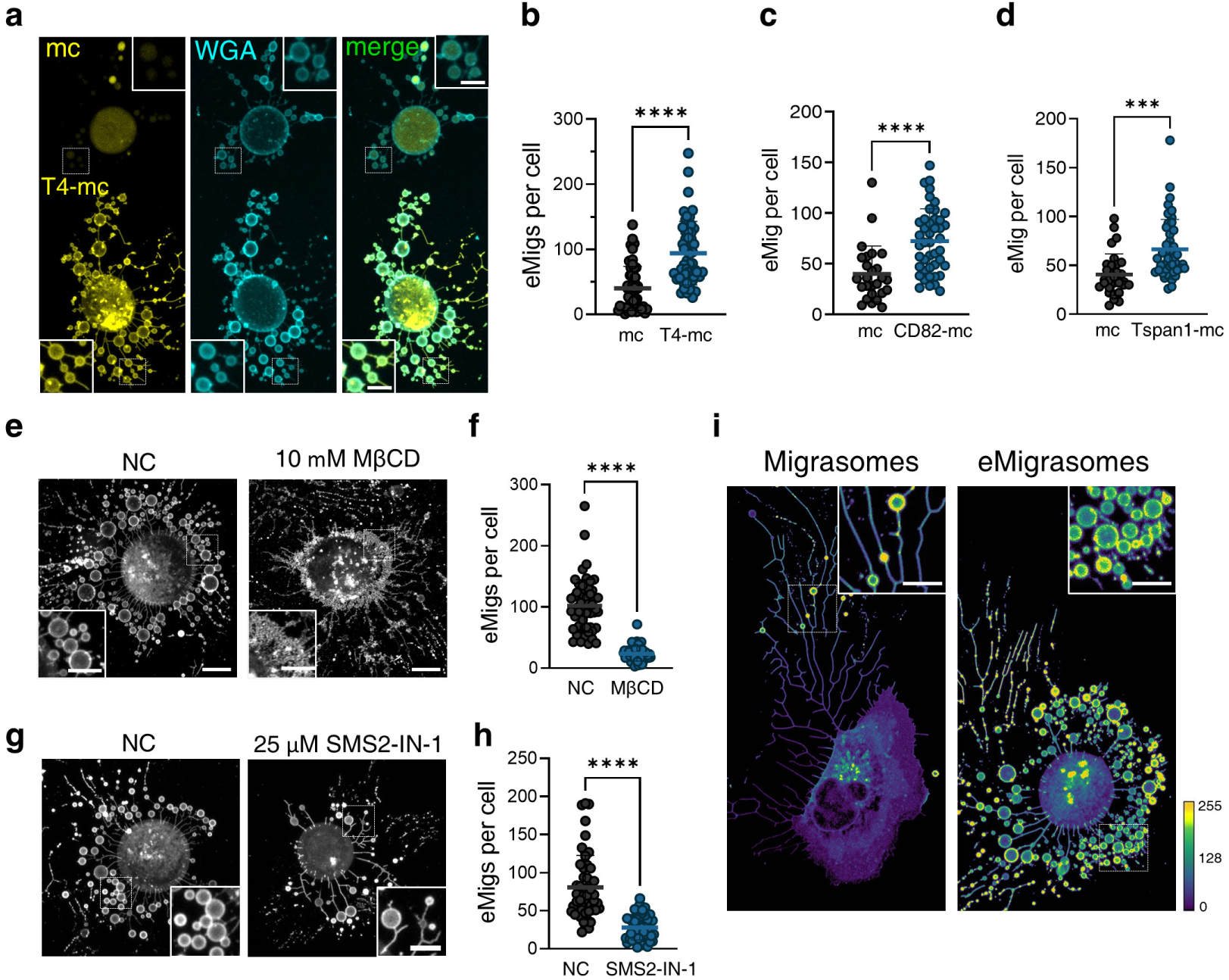
Mechanistic and morphologic similarity between migrasomes and eMigrasomes. (a) Representative confocal images showing the effect of Tspan4-GFP in the biogenesis of migrasome-like structures. NRK cells were transiently transfected with mCherry vector or Tspan4-mCherry. The two populations of transfected cells were mixed in a 1:1.5 ratio in a test tube and then seeded in a confocal chamber. Cells were pre-incubated with 2 μM LatA for 10 mins and then treated with a three-step hypotonic stimulation with 2 min intervals. In each step, the osmolarity was reduced by 1/6 (16.7%). WGA-AF647 (Thermo, W32466) was then added to stain migrasome-like structures. Z-stack images were captured for further analysis. Scale bar, 5 μm. (b) Statistical analysis of the number of migrasome-like structures per cell in NRK cells transiently transfected with mCherry vector or Tspan4-mCherry in (a). n = 53, 53 cells respectively. (c) Statistical analysis of the number of migrasome-like structures per cell in NRK cells transiently transfected with mCherry vector (n = 26 cells) or CD82-mCherry (n = 44 cells) in Fig S1a. (d) Statistical analysis of the number of migrasome-like structures per cell in NRK cells transiently transfected with mCherry vector (n = 31 cells) or Tspan1-mCherry (n = 44 cells) in Fig S1b. (e) Representative confocal images showing the effect of cholesterol extraction on migrasome-like structures. NRK cells stably expressing Tspan4-GFP were stimulated to generate migrasome-like structures as described in (a). Cells were then incubated with 10 mM MβCD or buffer supplied with an equal volume of control solvent (H_2_O) for 30 min before imaging. Z-stack images were captured for further analysis. Scale bar, 10 μm. Insert scale bar, 5 μm. (f) Statistical analysis of the number of migrasome-like structures per cell in control cells (n=56) or cells treated with 10 mM MβCD (n=58) in (e). (g) Representative confocal images showing the effect of sphingomyelin depletion on the biogenesis of eMigrasomes. NRK cells stably expressing Tspan4-GFP were incubated with DMSO or 25 μM SMS2-IN-1 for 16 hrs. Cells were then treated and imaged as described in (a). Scale bar, 5 μm (h) Statistical analysis of the number of migrasome-like structures per cell in control cells (n=51) or cells treated with 25 μM SMS2-IN-1 (n=46) in (g). (i) Representative confocal images showing a cell generating natural migrasomes (left) and eMigrasomes (right). Migrasomes and eMigrasomes are morphologically similar. The fluorescence signal of Tspan4-GFP is highly enriched in both migrasomes and eMigrasomes.

Because of the mechanistic similarity between migrasomes and migrasome-like structures (Fig 2i), and because of the artificial nature of the procedure to generate these structures, we named these migrasome-like structures as engineered migrasomes (eMigrasomes).

Based on these results, we designed a protocol to generate and isolate eMigrasomes (eMigs) (Fig 3a). In this protocol, cells cultured in flasks were pretreated with high-potassium DPBS (K-DPBS) containing LatA and then the osmolarity was reduced by adding water in a step-wise manner. The flask then was subjected to moderate rotation to separate poorly-adhering cell bodies and tightly-adhering eMigs. The supernatant containing the majority of cell bodies was discarded and the eMigs attached to the bottom were harvested by pipetting. No trypsinization was applied to ensure optimal protection of the integrity of eMigs. To remove remaining cell bodies, crude eMigs were subjected to differential centrifugation followed by a gravity-dependent filtration through a 6-μm filter. eMigs in the flowthrough were concentrated by high-speed centrifugation. This protocol generated eMigs in a high yield. By microscopic examination, every individual cell robustly generates eMigs (Supplementary Video 4). By confocal microscopy analysis, the isolated eMigrasomes are round vesicles with different sizes (Fig 3b). To measure the size of eMigrasomes, we carried out a confocal-based analysis using Hough circle transformation (Fig 3b). We found that eMigrasomes have a size range from 1.4 μm - 6.6 μm, with a median size of 1.6 μm (Fig 3c). Since negative staining might distort the shape of membrane vesicles (Fig 3d), we carried out cryo-EM analysis of eMigrasomes. Under cryo-EM, eMigrasomes appeared as intact round vesicles (Fig 3e). Notably, the eMigs were not contaminated with significant amounts of intracellular membranes, as western blotting of purified eMigs showed very little contamination from other organelles (Fig 3f).

**Figure 3.**
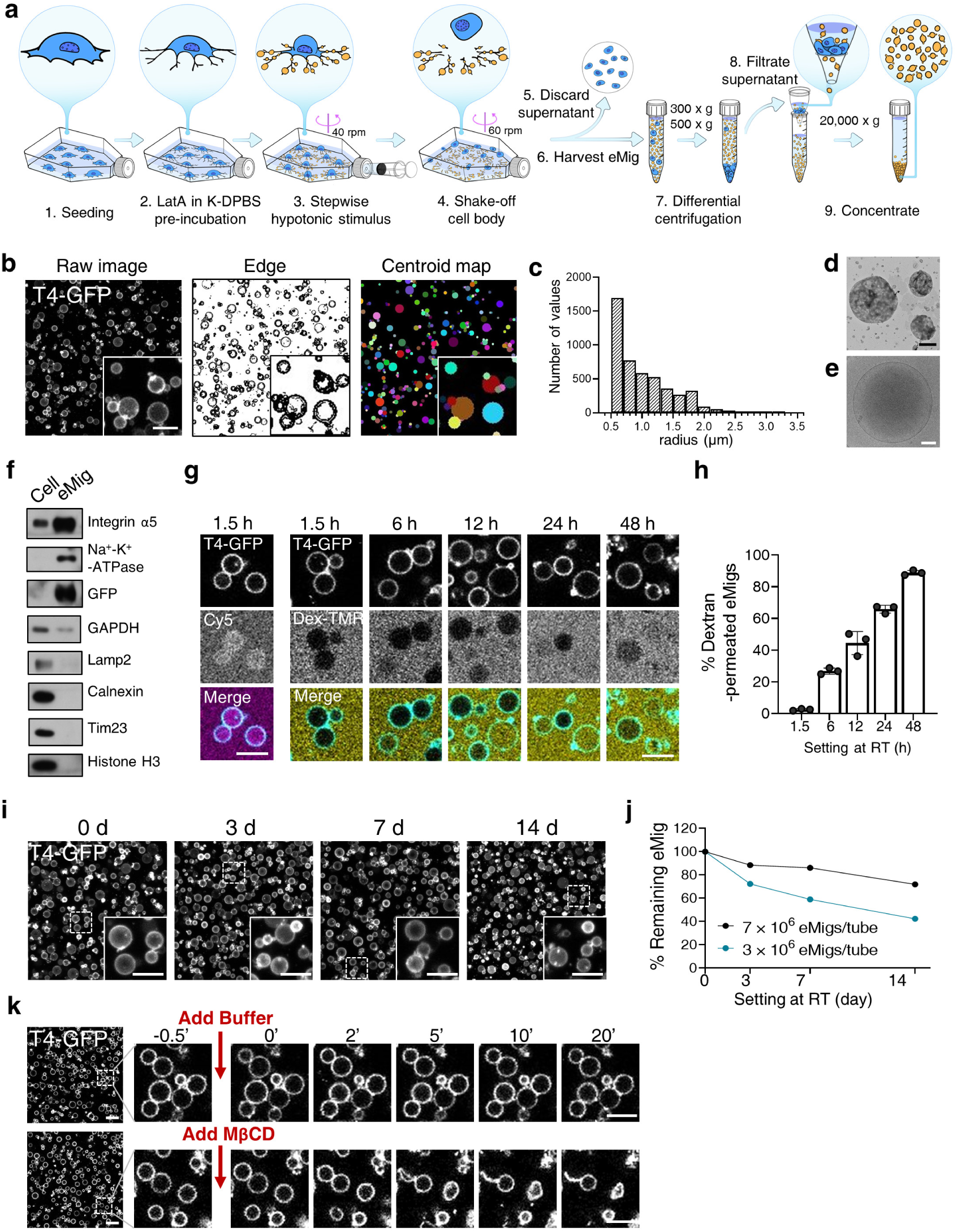
Isolation and characterization of eMigrasome. NRK cells stably expressing Tspan4-GFP were used in all experiments in this figure if not otherwise specified. (a) Schematic illustration showing the process of eMigrasome induction, isolation and purification. (b) Confocal image (left), threshold edge (middle) and centroid map (right) of purified eMigrasomes. Image processing and analysis were performed using imageJ. The Hough circle transform plugin was applied to recognize and transform thresholded edges into binned objects representing individual eMigrasomes. Scale bar, 5 μm (c) Statistical analysis of the radius of purified eMigrasomes. Measurement was performed using the map generated by Hough circle transformation analysis. 4725 particles were analyzed and the data were binned to plot the distribution of eMigrasomes radius. (d) TEM micrograph of negatively stained purified eMigrasomes. Scale bar, 1 μm. (e) Cryo-EM micrograph of purified eMigrasomes. Scale bar, 200 nm. (f) Western blot showing the protein level of several markers in cell bodies and eMigrasomes. An equal amount of protein was loaded in each lane. (g) Representative time-lapse confocal images showing the high permeability of eMigrasomes to Cy5 at 1.5 hrs post purification (left) and the gradual increase in the permeability of eMigrasomes to 40 kDa dextran-TMR (right). Scale bars, 5 μm (h) Statistical analysis of the percentage of eMigrasomes that were permeable to 40 kDa dextran-TMR at the indicated timepoints. For each time point, eMigrasomes from three different views were analyzed. From left to right, n = 336, 493, 812, 830 and 631. (i) Representative confocal images of eMigrasomes after sitting at room temperature for 0, 3, 7 or 14 days. Aliquots of eMigrasomes were stored in EP tubes as pellets at room temperature for the indicated time, then resuspended and dropped into a confocal chamber before imaging. Z-stack image series were captured and sum-slices projections were applied. Scale bar, 5 μm. (j) Number of eMigrasomes after storage at room temperature for 0, 3, 7 or 14 days. Aliquots of eMigrasomes (7 x 10^6^ eMigrasomes per tube in black, 3 x 10^6^ eMigrasomes per tube in green) were stored in EP tubes as pellets at room temperature for the indicated time, then resuspended and stained with WGA561 before counting by FACS. (k) Time-lapse image series showing purified eMigrasomes treated with 10 mM MβCD or control buffer. A 10 μl drop of concentrated eMigrasomes was settled in a confocal chamber, then sealed and maintained at 37°C during imaging. Buffer containing 10 mM MβCD or control solvent was added to the drop using the equipment illustrated in Fig 1d. Scale bar, 5 μm

Previously we reported that migrasomes can become leaky before rupture. We wondered whether eMigrasomes can also become leaky. To test this, we added eMigrasomes into isolation buffer loaded with Cy5 and 40 kDa dextran-TMR. At room temperature, eMigrasomes rapidly become leaky to Cy5, a fluorescent dye that does not pass intact membranes (Fig 3g). During the 48 hours after preparation, eMigrasomes gradually become leaky to 40 kDa dextran-TMR, which mimics the size of a normal protein (Fig 3g, 3h).

Next, we tested the stability of eMigrasomes at room temperature. Surprisingly, eMigrasomes are highly stable. Even after 14 days at room temperature, the morphology of eMigrasomes did not change significantly, and the number of eMigrasomes was only slightly reduced (Fig 3i, 3j). However, if we treated isolated eMigrasomes with MβCD, most of them deformed and ruptured within 30 min (Fig 3k). This suggests a crucial contribution of cholesterol to the stability of isolated eMigrasomes.

The stable nature of eMigs prompted us to explore the possibility of using eMigs as carriers for delivery of proteins (Fig 4a). We found that membrane proteins (e.g. the cell surface receptor PD1) can be easily loaded onto eMigrasomes by simply overexpressing these proteins in cells (Fig 4b). To load cytosolic proteins, we fused them with the transmembrane domain followed by the polybasic tail of syntaxin 2 (STX2). This allowed us to successfully load the cytosolic protein ovalbumin (OVA) onto the plasma membrane and thus onto eMigrasomes, referred as mOVA for membrane-tethered OVA (Fig 4c).

**Figure 4.**
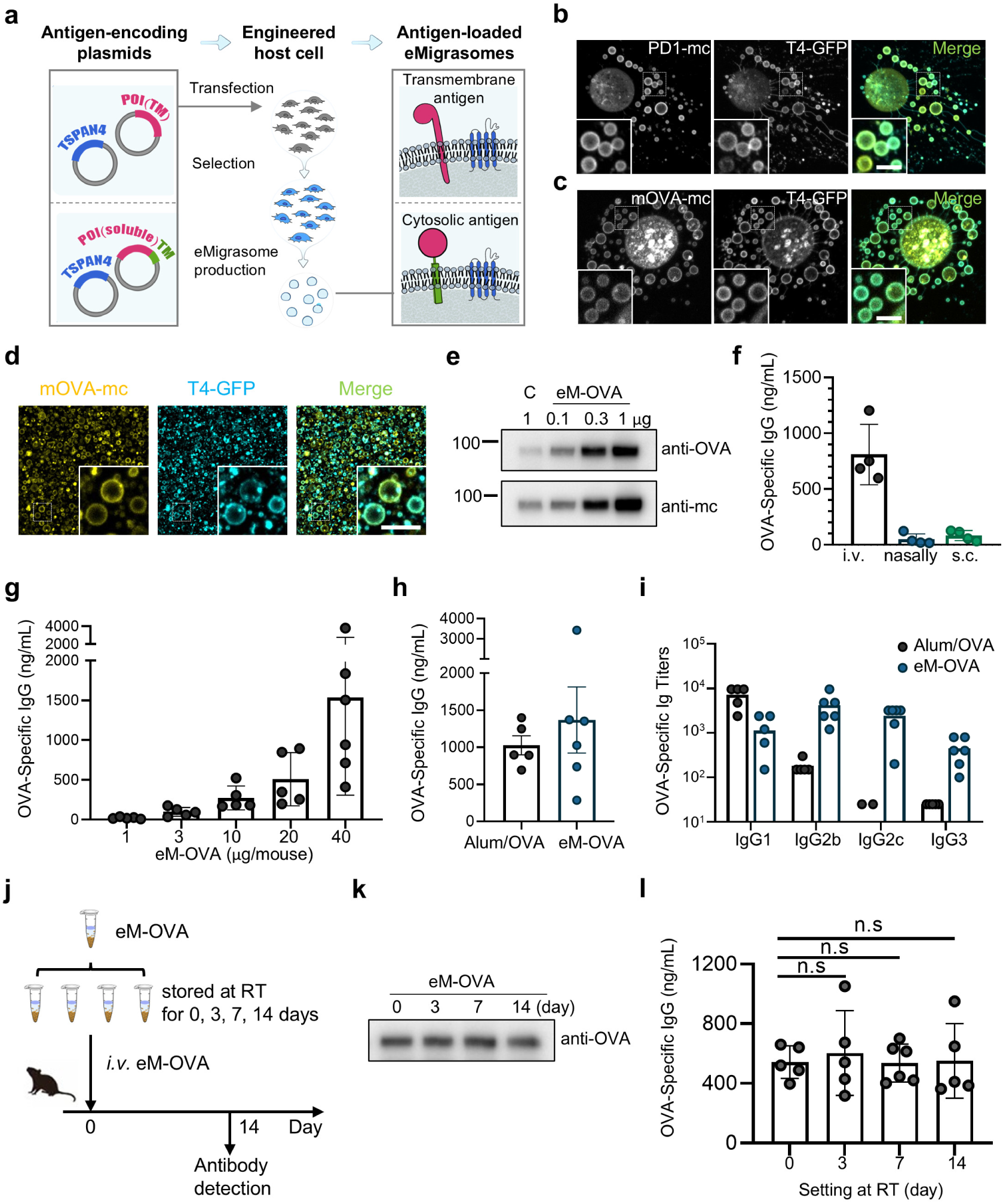
eMigrasomes as an antigen carrier for vaccination. (a) Schematic illustration of strategies for loading proteins of interest (POI) onto eMigrasomes. The protocol includes the construction of antigen-encoding plasmids, the construction of engineered cell lines stably expressing the antigens and the production of antigen-loaded eMigrasomes. Membrane proteins, which already carry a transmembrane (TM) domain, are overexpressed in cells (top). Cytosolic proteins are membrane tethered by tagging with the TM sequence and polybasic tail of STX2 (bottom). Cells are then treated with hypotonic buffer to enrich the POI on the surface of eMigrasomes. (b) Representative confocal image of a cell expressing PD1-mCherry and Tspan4-GFP. The transmembrane protein PD1-mCherry was loaded onto eMigrasomes as shown in the top part of (a). Scale bar, 5 μm (c) Representative confocal image of a cell expressing membrane-tethered OVA-mCherry (mOVA-mc) and Tspan4-GFP. To create an extracellular membrane-tethered form of OVA (mOVA), the sequence of OVA was fused to the C-terminus of a truncated form of mouse STX2, in which only the transmembrane region and a polybasic tail remained. The mCherry tag was fused to the C-terminus of OVA to trace the localization of this fusion protein. The soluble protein OVA was loaded onto the membrane of eMigrasomes, as shown in the bottom part of (a). Scale bar, 5 μm. (d) Representative confocal image of eMigrasomes isolated from MCA-205 cells stably expressing Tspan4-GFP and mOVA-mCherry (eM-OVA). Scale bar, 5 μm. (e) Western blot showing the amount of full-length mOVA-mCherry protein in host cell and eM-OVA. Cell lysate (C) containing 1 μg total protein and purified eM-OVA samples containing 0.1, 0.3 or 1 μg total protein were loaded. The protein-immobilized PVDF membrane was firstly incubated with anti-OVA antibody and then stripped and re-blotted with anti-mCherry antibody. The antigen mOVA-mCherry was highly enriched in eMigrasomes compared to host cells. MCA-205 cells stably expressing Tspan4-GFP and mOVA-mCherry were used for all experiments in the rest of this figure if not otherwise specified. (f) Amount of OVA-specific IgG in mouse serum on day 14 after intravenous (i.v), nasal or subcutaneous (s.c) immunization with eM-OVA (20 µg/mouse). OVA-specific IgG was quantified by ELISA. (g) ELISA quantification of OVA-specific IgG in sera from wild-type (WT) mice on day 14 after tail intravenous immunization with eM-OVA at the indicated dose. (h) ELISA quantification of OVA-specific IgG in sera from WT mice on day 14 after tail intravenous immunization with eM-OVA (20 µg/mouse) or intraperitoneal immunization with Alum/OVA. (i) Titer analysis of OVA-specific IgG1, IgG2b, IgG2c and IgG3 in the sera from mice immunized with eM-OVA (20 µg/mouse) or Alum/OVA. Serum samples were collected on day 14. Each dot represents an individual serum sample. (j) Illustration of the experimental setup for assaying the stability of eM-OVA. (k) Immunoblotting analysis of the amount of OVA protein in samples of eM-OVA which were left at room temperature for 0, 3, 7 or 14 days. 2 µg protein was loaded in each lane. (l) ELISA quantification of OVA-specific IgG in sera from WT mice on day 14 after tail intravenous immunization with eM-OVA stored at room temperature for 0 days (D0), 3 days (D3), 7 days (D7) or 14 days (D14). 20 µg eM-OVA was injected per mouse.

### An eMigrasome-based vaccine

Next, we explored the potential of eMigrasomes as an antigen carrier for vaccines. First, we employed OVA, a well-established model antigen, to evaluate eMigrasomes as a platform for antigen delivery. We used imaging and immunoblotting to confirm the presence of the mOVA-mCherry protein (Fig 4d, 4e). It is worth noting that the mOVA antigen was highly enriched in isolated eMigrasomes compared to the host cells (Fig 4e). We immunized mice with OVA-loaded eMigrasomes (eM-OVA) via different routes and found that intravenous injection resulted in the highest IgG antibody titer (Fig 4f). eM-OVA induced the antigen-specific IgG response in a dose-dependent manner (Fig 4g). The IgG response induced by eM-OVA at 20 μg/mouse was comparable to traditional Alum/OVA at 50 μg/mouse (Fig 4h). Together, these data suggest that eM-OVA elicits a strong IgG response compared to traditional alum-based immunization.

IgG is the most abundant immunoglobin in human and mouse, with four different subtypes: IgG1, IgG2, IgG3 and IgG4 in human, and IgG1, IgG2a/c, IgG2b, IgG3 in mouse^23^. Different IgG subtypes are highly conserved but each has its unique immunological functions, e.g. acting through different Fc-gamma receptors (FcγRs) or binding to complement^24^. Thus, we characterized the type of IgG induced by eM-OVA in mice, and compared it to that induced by Alum/OVA. The IgG response to Alum/OVA was dominated by IgG1, consistent with previous publications^25, 26^. Quite differently, eM-OVA induced an even distribution of IgG subtypes, including IgG1, IgG2b, IgG2c, and IgG3 (Fig 4i). Usually, the ratio between IgG1 and IgG2a/c indicates a Th1 or Th2 type humoral immune response. Thus, eM-OVA immunization induces a balance of Th1/Th2 immune responses.

Next, we assessed the stability of the immunogenicity of eM-OVA over a period of 14 days at room temperature. For this purpose, we placed the purified OVA-loaded eMigrasomes in a test tube at room temperature without adding any reagents to inhibit protein degradation. After 14 days of room temperature storage, the amount of intact OVA was roughly the same as in freshly purified eM-OVA, and the IgG response induced by eM-OVA remained unchanged during this period (Fig 4j – 4l).

### An eMigrasome-based vaccine induces a strong humoral protective response against SARS-CoV-2

We next explored eMigs as a platform to carry the SARS-CoV-2 Spike protein (S protein). To prevent the S protein from being broken down by proteases, we mutated the furin site of the S protein. MCA-205 cells were used to express the S protein with an mCherry tag. After isolating the eMigs, we used imaging to confirm the presence of the Spike protein (Fig 5a). To assess the antigen integrity, we performed immunoblotting using antibodies against both S1 and mCherry. Two distinct bands were observed: one at the expected molecular weight of the S-mCherry fusion protein, and a higher molecular weight band that may represent oligomerized or higher-order forms of the Spike protein (Fig 5b). Furthermore, we performed confocal microscopy using a monoclonal antibody against Spike. Co-localization analysis revealed strong overlap between the mCherry fluorescence and anti-Spike staining, confirming the proper presentation and surface localization of intact S-mCherry fusion protein on eMigs (Fig 5c). These results confirm the structural integrity and antigenic fidelity of the Spike protein expressed on eMigs.

**Figure 5.**
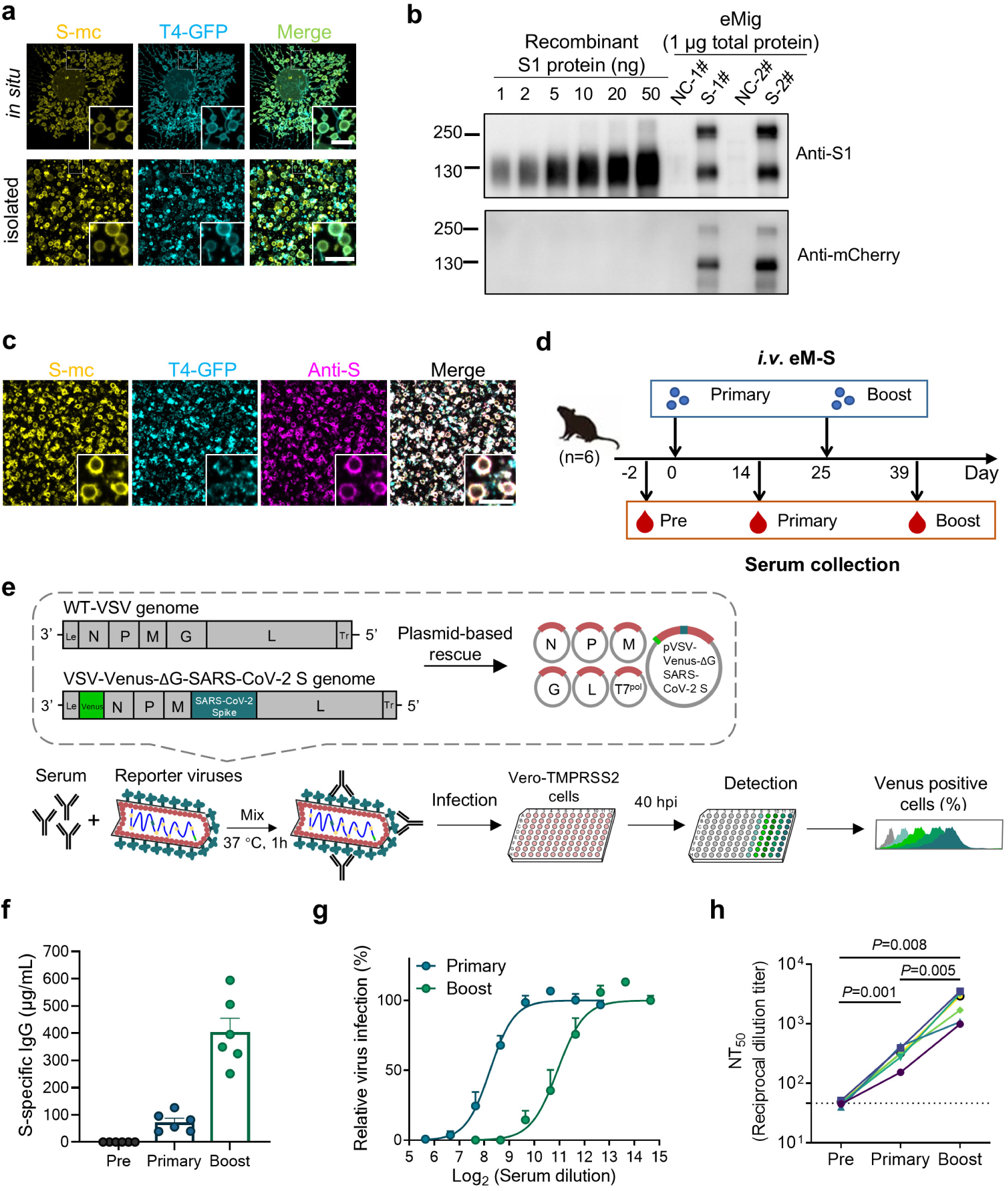
An eMigrasome-based vaccine induces a strong humoral protective response against SARS-CoV-2. (a) Representative confocal images showing the presence of Spike-mCherry (S-mc) in engineered cells and isolated eMigrasomes. Scale bar, 5 μm. (b) Western blot showing the amount of full-length Spike-mCherry protein in eM-S. A titration (1, 2, 5, 10, 20 or 50 ng) of recombinant spike protein were loaded as standards. Purified eM-NC or eM-S samples containing 1 μg total protein were loaded. (c) Representative confocal images showing the presence of integral spike protein in isolated eMigrasomes. Scale bar, 5 μm. (d) Schematic diagram of the experimental procedure for immunization with Spike-loaded eMigrasomes (eM-S) and collection of serum. (e) Illustration of the rVSV-venus-SARS-CoV-2 system. VSV-Venus-SARS-CoV2 was mixed with vaccinated mice sera. Vero-TMPRSS2 cells were infected with the reporter virus/serum mixture with an MOI of 0.01. 40 hrs post infection, the Venus-positive infected cells were quantified to estimate the NT_50_ value for each serum. (f) Spike-specific IgG was quantified in WT mice immunized with eM-S (20 µg/mouse, *i.v.*) at different time points. Each symbol represents one individual animal. (g) Neutralization curves are presented for sera from primary-vaccination and boost-vaccination. Dots along the lines represent means from individual serum samples. Nonlinear regression was performed using the equation for the normalized response versus the inhibitor, incorporating a variable slope. (h) NT_50_ of individual mouse in vaccinated groups were compared by p-value (Paired t-test) are indicated. Dotted lines represent assay limits of detection. Each line represents an individual mouse.

Immunization of WT C57BL/6J mice with the Spike-loaded eMigs (eM-S) resulted in a strong antibody response against the S protein, which was further improved by a second shot (Fig 5d, 5f). To assess the neutralizing capacity of the antisera provoked by eM-S, we utilized a replication-competent, infectious VSV chimera incorporated with the SARS-CoV-2 spike protein for a neutralization test (Fig 5e), similar to the previously reported system^27, 28^. This genetically altered VSV chimera virus features the SARS-CoV-2 spike protein, substituting its native surface glycoprotein (G), making the VSV chimera reliant on the SARS-CoV-2 spike protein for cellular entry. The Venus-based fluorescence reporter system offers high sensitivity. The neutralizing power of antisera against the SARS-CoV-2 spike protein was assessed by calculating the percentage of Venus-positive infected cells when treated with serum versus mock controls. The antisera, stimulated by an initial immunization with eM-S, neutralized the recombinant VSV-Venus-SARS-CoV2 up to a dilution titer of approximately 300 to achieve 50% neutralization (NT_50_). A secondary booster immunization further enhanced the neutralization capability with an NT_50_ up to 4000 (Fig 5g, 5h). Together, these results indicate that antigen-carrying eMigs can induce a strong humoral protective response against SARS-CoV-2.

## Discussion

In this manuscript, we describe a method to rapidly generate eMigrasomes from cultured mammalian cells. We developed a simple method to load membrane or cytosolic proteins onto eMigrasome. Using OVA as a model antigen, we demonstrate that eMigrasomes are a highly effective, temperature-stable vaccine platform which can elicit antibody response with a very small amount of antigen. Finally, we show that eMigrasomes can be used to generate effective vaccines against SARS-CoV-2. Collectively, our study provides the proof of concept for developing eMigrasome-based vaccines.

Previously, migrasomes have been defined as “migration-dependent” vesicles. In this study, we demonstrated that migrasome-like structures can be induced through the relative movement of the cell edge in a migration-independent manner. Notably, eMigrasomes exhibit conserved genetic and morphological features compared to natural migrasomes, providing strong evidence for this concept. Accordant with our study, a recent investigation has revealed that cell shrinkage induced by bacterial toxins can also trigger migrasome formation^29^.

In this study, we demonstrate the efficiency of eMigrasome-based vaccines using a model antigen or an antigen from a coronavirus. The benefit of using a model antigen is that there are readily available experimental systems. The benefit of using a well-characterized virus antigen is that it allows us to reliably test the efficacy of our vaccine platform in a realistic setting. It should be noted that, both a signal peptide and a transmembrane domain are necessary for proper antigen presentation on eMigrasomes. For antigens that do not naturally contain these two features, a non-native signal peptide or an artificial transmembrane domain should be engineered into the coding sequence of the antigen. We are aware that due to the experimental nature of this work, it is unlikely that the eMigrasome platform will be used to generate mainstream preventive vaccines in the foreseeable future. However, there is a very real possibility of developing eMigrasomes into therapeutic vaccines against cancer or other diseases with unmet medical needs, or into preventive vaccines against pathogens for which the current vaccine platforms do not work. Our investigation regarding the biophysical and cellular mechanisms of eMigrasome formation makes it possible for further optimizing eMigrasome generation in a rational manner.

In this study, we demonstrate that eMigrasomes can be an effective vaccine platform. In addition, we speculate that eMigrasomes may also emerge as a versatile delivery system for a range of applications. The high rigidity and self-repair capacity of migrasomes, which is caused by enrichment of tetraspanins and cholesterol, makes migrasomes highly stable, and thus suitable as delivery carriers in various *in vivo* settings. More importantly, the physiological roles of migrasomes indicate that they have naturally evolved as carriers for delivering materials and information *in vivo*. Our previous work showed that during embryonic development, migrasomes which are highly enriched with signaling molecules, such as chemokines, growth factors and morphogens, are deposited at various spatially defined locations, where they act as sustained-release capsules to liberate the signaling molecules. In this way, migrasomes affect multiple aspects of embryonic development including organ morphogenesis and angiogenesis ^30, 31^. Additionally, in cultured fibroblast cells, migrasomes are enriched with a selected set of full-length, translationally competent mRNAs. When these mRNA-enriched migrasomes are taken up by neighboring cells, the mRNA can escape the endo-lysosome system of recipient cells and be translated into protein, thus modifying the behaviors of the recipient cells ^32^. Finally, in cells experiencing mild mitochondrial damage, the damaged mitochondria can be selectively transported into migrasomes and then evicted from the cell in a process named as mitocytosis ^33^. In summary, multiple types of cargos, including materials and information, can be enriched in migrasomes under different settings, and migrasomes can be deposited at spatially defined locations in diverse biological settings to affect a broad range of biological processes, including cell-cell communication. Since eMigrasomes capture certain key features of migrasomes, it is our speculation that we may able to develop eMigrasomes into a delivery system for diverse cargo types including nucleic acids, proteins, small molecules and even organelles.

## Supplementary Information

**Figure S1.**
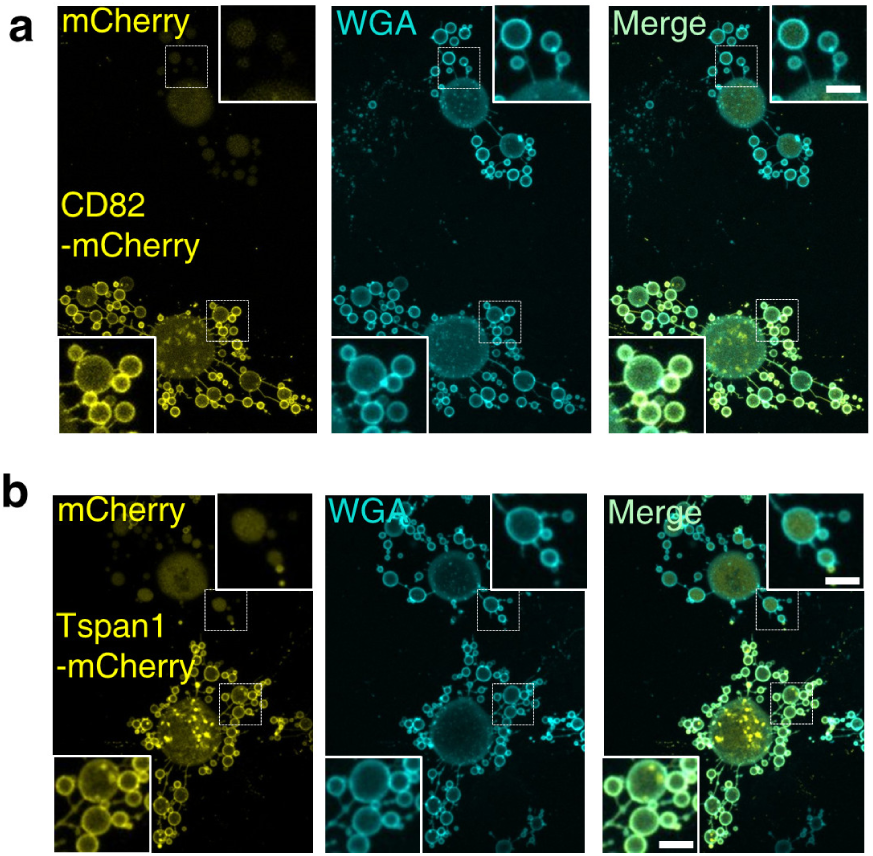
The effect of tetraspanin expression on the formation of migrasome-like structures. (a) NRK cells were transiently transfected with mCherry vector or CD82-mCherry. The cells were then treated and imaged as described in Fig 2a. (b) NRK cells were transiently transfected with mCherry vector or Tspan1-mCherry. The cells were then treated and imaged as described in Fig 2a.

## Supplementary video legends

**Supplementary video 1**

Time-lapse movie showing the biogenesis of migrasome-like structures. Related to Figure 1a.

**Supplementary video 2**

The dynamic of migrasome-like structures induced by single-step approach. Related to the upper panel of Figure 1e.

**Supplementary video 3**

The dynamic of migrasome-like structures induced by stepwise approach. Related to the lower panel of Figure 1e.

**Supplementary video 4**

4D time-lapse movie showing the biogenesis of eMigrasomes.

## Materials and Methods

### Molecular cloning

The sequence spanning amino acids 252 to 289 of syntatin-2 (STX-2), which incorporates the transmembrane domain along with the polybasic tail, was amplified from mouse cDNA by polymerase chain reaction (NEB, M0492L) using primers below:

fwd:5’- atgaagaaagccatcaaataccagagc −3’;

rev:5’- gcaccgatggagcccatcgagcctttgccaaccgacaagccaatg −3’.

The sequence of ovalbumin was amplified from the plasmid using primers below:

fwd: 5’- atgggctccatcggtgc −3’;

rev: 5’- aggggaaacacatctgccaaag −3’.

Using STX2_252-289_ and ovalbumin as templates, the membrane-tethered mOVA, in which STX2_252-289_ sequence is tagged to ovalbumin sequence, was further amplified using the primers below:

fwd: 5’- tcagatctcgagctcaagcttatgaagaaagccatcaaataccagagc −3’

rev: 5’- tggtggcgaccggtggatcccggaggaagaacactaaggcagcaaaagagaag −3’

The fragment was inserted into pmCherry-N1 vector using a One Step Cloning Kit (Vazyme, C112).

### Cell culture

Cells were cultured at 37 °C with 5% CO_2_. NRK cells and HEK 293FT cells were grown in DMEM (Gibco, C11995500BT) supplemented with 10% (v/v) FBS (Biological Industries), 1% (v/v) glutamax (Gibco, 35050-061) and 1% (v/v) penicillin– streptomycin. MCA-205 cells were grown in RPMI 1640 (Gibco, C11875500BT) supplemented with 10% (v/v) FBS, 1% (v/v) glutamax and 1% (v/v) penicillin– streptomycin Vero-TMPRSS2 cells were maintained in DMEM supplemented with 10% (v/v) FBS and 50 IU/ml penicillin-streptomycin. ADSC cells and BMSC cells were grown in stem cell serum-free medium (CytoNiche, RMZ112).

### Cell transfection and cell line development

For NRK cells, transfection was performed using electroporation (AMAXA, Nucleofector). For MCA-205 cells, transfection was performed using Lipofectamine 3000 transfection reagent (Thermo Fisher Scientific, L3000015). In establish stable cell lines, the transfected cells underwent initial selection with Hygromycin B (Roche, 10843555001) and were subsequently sorted into single colonies in 96-well plates via flow cytometry.

### Imaging

Confocal imaging of live cells and isolated eMigrasomes was performed using a Nikon A1HD25 laser scanning confocal microscope. Cells were cultured in a confocal chamber (Cellvis, D35-20-1-N or D35C4-20-1.5-N) that had been pre-treated with 10 μg/ml fibronectin (F0895) and allowed to proliferate overnight (14-18 hours). For visualizing eMigrasomes, a small region in the confocal chamber was coated with a 10 μl drop of 10 μg/ml fibronectin. Following aspiration of the fibronectin, a 10 μl drop of isolated eMigrasomes was dispensed into the coated area and left to settle for a minimum of 1 hour prior to imaging. To prevent evaporation, the chamber was sealed using parafilm. For immunofloresence assay, eMigrasomes were settled at 4 °C overnight and then fixed by 2% PFA. Nonspecific binding was blocked by incubating with 10% (v/v) FBS. eMigrasomes were then stained with a spike primary antibody (40150-D001, SinoBiological) for 1 hours at RT, triple washed with PBS and then stained with AlexaFlour647-conjugated secondary antibody. Samples were triple washed with PBS before imaging.

### Real-time hypotonic stimulation

To observe the dynamics of migrasome-like structures and eMigrasomes, a custom-made buffer displacement device was utilized (refer to Fig 1d). Essentially, three apertures were created on the lid of a confocal chamber to perfectly accommodate a silicon microinjection tube with an external diameter of 1.9 mm. The silicon microinjection tube was securely connected to a 1 ml syringe after the removal of its sharp needle.

For a single-step stimulation process, two syringes were employed. One was kept empty, while the other was filled with the desired hypotonic buffer. During time-lapse imaging, the reservoir solution was aspirated using the empty syringe, and the hypotonic buffer was simultaneously injected from the other syringe.

In the case of stepwise stimulation, three syringes were deployed. The first was loaded with the initial buffer from the reservoir and was used to withdraw the reservoir solution. The second syringe contained deionized water (ddH_2_O) and was used to add a specific volume of water at each step. The third syringe, which was empty, was used to ensure thorough mixing of the reservoir solution after each stimulation event.

### Drug treatment

For SMS2 inhibition, 30 min post seeding of the cells, the culture medium was gently replaced with fresh medium containing either 25 μM SMS2-IN-1 (MCE, HY-102041) or 0.25 (v/v) % DMSO. Cells were treated for 16 hrs prior to imaging.

For cholesterol extraction from *in situ* or isolated eMigrasomes, h-KDPBS containing 10 mM MβCD (Sigma, 332615) or 10 (v/v) % ddH_2_O was applied. For *in situ* eMigrasomes, z-stack images were collected after 30 min incubation at 37 °C. For isolated eMigrasomes, buffer containing MβCD or ddH_2_O was applied using the real-time stimulation device described above. A 20-min time-lapse imaging session was performed to capture the process of morphological changes in eMigrasomes.

### Permeability assay

Isolated eMigrasomes were diluted in h-KPBS-BSA, containing 5 μg/ml of Cy5 and 25 μg/ml of 40kDa Dextran-TMR (D1842). A droplet of the eMigrasome solution was prepared and imaged following the procedure described previously.

### Ion replacement assay

The standard formulation of DPBS is 138 mM NaCl, 8.1 mM Na_2_HPO_4_, 2.7 mM KCl, 1.5 mM KH_2_PO_4_. In the ion replacement assay, as referenced in Figure S1d and S1e, the 138 mM NaCl in DPBS was substituted with equal molar KCl, CsCl or Choline chloride. The other three components were not changed.

The K-DPBS formulation utilized for eMigrasome production contains 140.6 mM KCl, 1.5 mM KH_2_PO_4_, 8.1 mM K_2_HPO_4._

### Induction and purification eMigrasomes

It’s important to underscore that several critical parameters, such as the seeding density, the working concentration of latrunculin A, and the incubation period, along with the hypotonic gradient, should be individually optimized for each cell type. The quantities of cells and reagents utilized should be modulated according to the flask size (examples are provided in the following table).

| Cell type | NRK | MCA-205 | ADCS | BMSC |
| --- | --- | --- | --- | --- |
| Seeding density (cells / cm<sup>2</sup>) | 1.5 x 10<sup>4</sup> | 3 x 10<sup>4</sup> | 2 x 10<sup>4</sup> | 2 x 10<sup>4</sup> |
| Latrunculin A concentration (μM) | 2 | 2 | 0.6 | 0.6 |
| Latrunculin A incubation time (min) | 10 | 45 | 20 | 20 |
| Reduced osmolarity per step | 1/6 | 1/4 | 1/6 | 1/6 |

In the following method, the purification of eMigrasomes from NRK cells cultured in a single T75 flask (NEST, 708003) is used as a representative example. Three major steps include cell preparation, eMigrasome induction and purification.

I. Cell Preparation

A T75 flask was coated with 4 μg/ml of fibronectin. 1×10^6^ Cells were seeded into the pre-coated flask, allowed to grow for 14-16 hours.

II. eMigrasome induction

Cortical actin disruption was achieved by discarding the culture medium and rinsing the cells once with PBS. Then, 7.5 ml of K-DPBS containing 2 μM of Latrunculin-A was added to the flask. The cells were incubated at 37°C in a 5% CO_2_ atmosphere for 10 minutes.

Hypotonic stimulation was performed by placing the flask on an orbital shaker inside the CO_2_ incubator and adjusting the speed to 40 rpm. The stimulation was progressively introduced by adding 1.5 ml, 1.8 ml, or 2.2 ml of deionized water at three-minute intervals. Following this, the speed of the orbital shaker was increased to 60 rpm for five minutes.

III. eMigrasome purification

First, the bubbles were harvested using the following procedure. The supernatant containing cell bodies was discarded. The flask’s bottom was gently washed once with hK-DPBS (60% K-DPBS matching the current osmolarity). Subsequently, 4 ml of hK-DPBS-BSA (60% K-DPBS containing 1 mg/ml BSA) was added to the flask. Gentle pipetting was employed to detach the eMigrasomes, and the resultant solution was collected into a conical tube. This step was repeated once, and the collected solutions were combined.

To remove the remaining cell bodies, a process of differential centrifugation and gravity-dependent filtration was used. Initially, the solution was centrifuged at 300 x g at 4°C for 10 minutes, after which the supernatant was collected. The same process was repeated at 500 x g for 10 minutes. The collected supernatant was poured into a filter cup containing a pre-rinsed 6 μm parylene filter (Hangzhou Branemagic Medical

Technology, F-PAC007). The filter was pre-rinsed with 100% ethanol followed by hK-DPBS-BSA. The flow-through was then collected in a low-protein-binding conical tube (Eppendorf, 0030122240).

To concentrate the eMigrasomes, the solution was centrifuged at 20,000 x g at 4°C for 30 minutes. In the event of a large volume, a longer time may be necessary. The supernatant was discarded, and the eMigrasome pellet was resuspended in PBS. The protein concentration was determined using a BCA analysis, and the PBS volume was adjusted to reach the desired eMigrasome concentration. For long-term storage, it was recommended that the eMigrasomes were stored as a pellet, with a small volume of hK-DPBS-BSA to cover it. This measure was adopted to minimize eMigrasome loss due to container adsorption.

### Negative staining TEM

Isolated eMigrasomes underwent fixation by the addition of an equal volume of 2.5% glutaraldehyde (GA). A droplet of the fixed eMigrasome sample was deposited onto a copper grid for 15 minutes. Excess sample was blotted off with filter paper. The grid was promptly washed with a droplet of double distilled water (ddH_2_O), stained with 1% uranyl acid for one minute, and then further washed with two droplets of ddH_2_O. Remaining water was blotted off using filter paper and the grid was allowed to air-dry fully before imaging was conducted via transmission electron microscopy (TEM).

### Cryo-EM sample preparation and image acquisition

Quantifoil Cu grids (200 mesh, R2/2) underwent glow-discharging using a plasma cleaning device (PDC-32G, Harrick Plasma). Each EM grid was applied with 4 µl sample and vitrified by plugging into liquid ethane using the Vitrobot Mark IV system (Thermo Fisher Scientific). The cryo-EM samples were examined using an FEI Tecnai Arctica 200 kV transmission electron microscope, and images were captured at a magnification of 23.5 kx using an FEI Falcon II direct electron detector, with a dose approximately around 15 e−Å−2.

### eMigrasome quantification using flow cytometry

Isolated eMigrasomes were suspended in h-KDPBS containing 1 μg/ml Wheat Germ Agglutinin (WGA, Thermo, W11262). Various dilutions were prepared to ensure at least one dilution had a concentration ranging from 1000 to 10000 eMigrasomes per μl. Data were collected using a CytoFlex LX cytometer (Beckman). The threshold for forward scatter (FSC) was manually set to 4000 to detect small particles. Events that were double positive for B525-FITC and Y610-mcherry were gated as eMigrasomes.

### Size analysis by Hough Circle Transforming

Size analysis of isolated eMigrasomes was conducted using ImageJ, complemented by the Hough Circle Transform plugin. Sum slices processing of Z-stack was applied to the 488 channel of the confocal images, representing the fluorescence signal from Tspan4-GFP. The image was subsequently converted to an 8-bit grayscale. Edge detection was performed using the “Find Edges” function to identify individual eMigrasomes. Subsequently, the “Threshold” function was applied to encompass the majority of the fluorescent signal from the eMigrasomes. Hough Circle Transform analysis was then utilized with the following parameters: Easy mode; Minimum = 3 pixels; Maximum = 24 pixels; Hough score threshold = 0.9. All output options were selected.

### Immunoblotting

Cells or eMigrasomes were subjected to lysis using a 2% SDS solution in 50 mM Tris buffer, followed by heating at 95 °C. The protein concentration of the samples was assessed using a BCA kit (Vazyme, E112-02-AB). The lysates were diluted to the desired concentration and then denatured via the addition of loading buffer (Beyotime, P0015). The proteins were segregated by SDS-PAGE electrophoresis (Epizyme, PG112 or Yeasen, 36255ES10), and subsequently transferred to a 0.45 μm PVDF membrane (Millipore, IPVH00010) in accordance with a standard protocol. The blot was blocked with 5% milk in TBST and left to incubate overnight with the primary antibody at 4 °C. The blot was then washed thrice with TBST and incubated with the secondary antibody at room temperature for 1 hour. Finally, the blot underwent three more washes prior to signal detection via chemiluminescence imaging (CYANAGEN, XLS070P or Thermo, 34075), utilizing a ChemiDoc MP Imaging System (Biorad).

Primary antibodies used for immunoblotting included: anti-integrin α5 (CST, 4705T), anti-Na-K-ATPase (CST, 3010S), anti-histone H3 (CST, 4499S), anti-lamp2 (Sigma), anti-calnexin (abcam, ab22595). anti-GFP (Roche, 11814460001 or abcam, ab290), anti-GAPDH (Proteintech, 60004-1-Ig), anti-TIM23 (BD, 611222), anti-S1 (Sinobio, 40591-MM42), anti-ovalbumin (santa cruz, sc80587 or abcam, ab181688), anti-mCherry (abcam, ab125096). Primary antibodies were diluted using Solution 1 (Takara, NKB-101). Secondary antibodies used for immunoblotting included peroxidase AffiniPure Goat Anti-Rabbit IgG (H+L) (Jackson ImmunoResearch, 111-035-003) and peroxidase AffiniPure Goat Anti-Mouse IgG (H+L) (Jackson ImmunoResearch, 115-035-003). Secondary antibodies were diluted using 5% milk in TBST.

### Mice

WT C57BL/6 (Jax 000664) specific-pathogen-free (SPF) mice were procured from the Laboratory Animal Center of Tsinghua University, China. The OT-II (Jax 004194) mice were donated by Dr. Yan Shi. All the mice were bred and maintained under SPF conditions at the Laboratory Animal Center of Tsinghua University, in accordance with the National Institute of Health Guide for the Care and Use of Laboratory Animals.

### Immunization

For systemic immunization, mice were intravenously administered eM-OVA dissolved in 100 µL PBS, or intraperitoneally administered a mixture containing 50 μg OVA in 50 μL PBS (Sigma; A5503) and 50 μL Imject® Alum (Thermo; 77161). For nasal immunization, mice were anesthetized using isoflurane and subsequently intranasally administered 20 μg eM-OVA dissolved in 30 μL PBS. For subcutaneous immunization, 20 μg eM-OVA in 100 µL PBS was administered through injections on both sides of the buttock.

### Enzyme linked immunosorbent assay (ELISA)

Serum was prepared from whole blood by centrifugation. The levels of antigen-specific antibody were determined using a direct ELISA method. Briefly, a 96-well plate were coated overnight at 4 °C with antigen (2 μg/mL). The wells were then blocked with PBS containing 10 % fetal calf serum before the addition of serially diluted serum.

Horseradish peroxidase-conjugated secondary antibodies were incubated for 1 hour at room temperature. Between each step, wells were washed with PBST. The colorimetric reaction was carried out using the 1-Step Ultra TMB-ELISA Substrate. The reaction was stopped with 2 N H_2_SO_4_, and absorbance at 450 nm was read using a multimode reader. For quantification of OVA-IgG and Spike-IgG, a standard curve was generated using serially diluted anti-OVA antibody (Santa Cruz; sc-80589) and anti-Spike antibody (Sino Biological; 40591-MM42), respectively.

### Fluorescence-based neutralization test

Mouse sera were heat-inactivated at 56°C for a duration of 30 min. The indicated dilutions of samples were mixed with 2×10^2^ FFU (Focus Forming Units) of VSV-Venus-SARS-CoV-2 and incubated for 1 h at a temperature of 37°C. The mixture of serum and virus were then added to Vero-TMPRSS2 cells grown on 96-well plates, and incubated at 37°C for about 40 hours. The cells were then harvested and fixed in a 4% paraformaldehyde solution for 20 min at room temperature. The fixed cells were resuspended in PBS and analyzed using a LSRFortessa SORP (BD Biosciences) and FlowJo software.

The Vero-TMPRSS2 cell line, which was constructed based on Vero (ATCC CCL-81™), stably expressed the TMPRSS2 (Transmembrane Serine Protease 2) protein to enhance the entry of the SARS-CoV-2 spike.

### Statistical analysis

All data were subjected to analysis using GraphPad Prism statistical software. Unpaired two-tailed t-tests or paired two-tailed t-tests were employed for the data analysis. The results are represented as the mean ± standard error of the mean (s.e.m.). A *P* value of less than 0.05 was deemed to indicate statistical significance.

## Acknowledgements

This research was supported by the National Key Research and Development Program of China (2018YFE0207300), National Natural Science Foundation of China (grant no. 32030023), Beijing Municipal Science & Technology Commission, Administrative Commission of Zhongguancun Science Park (grant no. Z221100003422012), Tsinghua University Initiative Scientific Research Program (grant no. 20221080007). Sincere gratitude is expressed towards Baidong Hou (Institute of Biophysics, Chinese Academy of Sciences) for providing mouse strains and engaging in valuable discussions. We thank the State Key Laboratory of Membrane Biology, SLSTU-Nikon Biological Imaging Center, Laboratory of Animal Resources Center (THU-LARC), Tsinghua University, for technical support. Acknowledgment is also given to the Research Fund of the Vanke School of Public Health at Tsinghua University.

## Author contributions

L.Y. and Z.L. supervised the project. The experiments were designed by L.Y., Z.L., Q.D., D.W., H.W., and Z.Z. D.W. carried out all the experiments related to the discovery, development, and characterization of eMigrasomes. H.W. performed all the animal experiments and immunological analyses. Z.Z. conducted pseudovirus neutralization assays. W.W. participated in the immunological experiments. Takami. S. and Y.Z. provided assistance with molecular biology and cell biology experiments. X.Z. was responsible for collecting EM data. L.D. contributed to the purification of eMigrasomes. The manuscript was written by L.Y. and Z.L., with input from all the authors.

## References

1. Park, J.W., Lagniton, P.N.P., Liu, Y. & Xu, R.H. mRNA vaccines for COVID-19: what, why and how. International journal of biological sciences 17, 1446–1460 (2021).

2. Polack, F.P. et al. Safety and Efficacy of the BNT162b2 mRNA Covid-19 Vaccine. N Engl J Med 383, 2603–2615 (2020).

3. Baden, L.R. et al. Efficacy and Safety of the mRNA-1273 SARS-CoV-2 Vaccine. N Engl J Med 384, 403–416 (2021).

4. Gui, Y. et al. Safety and immunogenicity of a modified COVID-19 mRNA vaccine, SYS6006, as a fourth-dose booster following three doses of inactivated vaccines in healthy adults: an open-labeled Phase 1 trial. Life Metabolism 2, load019 (2023).

5. Crommelin, D.J.A., Anchordoquy, T.J., Volkin, D.B., Jiskoot, W. & Mastrobattista, E. Addressing the Cold Reality of mRNA Vaccine Stability. Journal of pharmaceutical sciences 110, 997–1001 (2021).

6. Grau, S., Ferrández, O., Martín-García, E. & Maldonado, R. Reconstituted mRNA COVID-19 vaccines may maintain stability after continuous movement. Clinical microbiology and infection : the official publication of the European Society of Clinical Microbiology and Infectious Diseases 27, 1698.e1691–1698.e1694 (2021).

7. Ma, L. et al. Discovery of the migrasome, an organelle mediating release of cytoplasmic contents during cell migration. Cell Res 25, 24–38 (2015).

8. Wu, D. et al. Pairing of integrins with ECM proteins determines migrasome formation. Cell Res 27, 1397–1400 (2017).

9. Huang, Y. et al. Migrasome formation is mediated by assembly of micron-scale tetraspanin macrodomains. Nat Cell Biol 21, 991–1002 (2019).

10. Ding, T. et al. The phosphatidylinositol (4,5)-bisphosphate-Rab35 axis regulates migrasome formation. Cell Res (2023).

11. Dharan, R. et al. Tetraspanin 4 stabilizes membrane swellings and facilitates their maturation into migrasomes. Nat Commun 14, 1037 (2023).

12. Huang, Y., Zhang, X., Wang, H.W. & Yu, L. Assembly of Tetraspanin-enriched macrodomains contains membrane damage to facilitate repair. Nat Cell Biol 24, 825–832 (2022).

13. Levy, S. & Shoham, T. The tetraspanin web modulates immune-signalling complexes. Nat Rev Immunol 5, 136–148 (2005).

14. Charrin, S. et al. EWI-2 is a new component of the tetraspanin web in hepatocytes and lymphoid cells. Biochemical Journal 373, 409–421 (2003).

15. Clark, K.L., Zeng, Z.-M., Langford, A.L., Bowen, S.M. & Todd, S.C. PGRL Is a Major CD81-Associated Protein on Lymphocytes and Distinguishes a New Family of Cell Surface Proteins1. The Journal of Immunology 167, 5115 - 5121 (2001).

16. Lefranc, M.-P. & Lefranc, G. The Immunoglobulin FactsBook. (2001).

17. Zucker, B. et al. Migrasome formation is initiated preferentially in tubular junctions by alternations of membrane tension or intracellular pressure. bioRxiv, 2023.2008.2003.551756 (2023).

18. Yoshikawa, K., Saito, S., Kadonosono, T., Tanaka, M. & Okochi, M. Osmotic stress induces the formation of migrasome-like vesicles. FEBS Letters (2024).

19. Hoffmann, E.K., Lambert, I.H. & Pedersen, S.F. Physiology of cell volume regulation in vertebrates. Physiol Rev 89, 193–277 (2009).

20. Qiu, Z. et al. SWELL1, a plasma membrane protein, is an essential component of volume-regulated anion channel. Cell 157, 447–458 (2014).

21. Voss, F.K. et al. Identification of LRRC8 Heteromers as an Essential Component of the Volume-Regulated Anion Channel VRAC. Science 344, 634–638 (2014).

22. Chen, L., Ma, L. & Yu, L. WGA is a probe for migrasomes. Cell Discov 5, 13 (2019).

23. Martin, R.M., Brady, J.L. & Lew, A.M. The need for IgG2c specific antiserum when isotyping antibodies from C57BL/6 and NOD mice. Journal of Immunological Methods 212, 187–192 (1998).

24. Vidarsson, G., Dekkers, G. & Rispens, T. IgG Subclasses and Allotypes: From Structure to Effector Functions. Frontiers in Immunology Volume 5 - 2014 (2014).

25. Comoy, E.E., Capron, A. & Thyphronitis, G. In vivo induction of type 1 and 2 immune responses against protein antigens. International Immunology 9, 523–531 (1997).

26. HogenEsch, H. Mechanism of Immunopotentiation and Safety of Aluminum Adjuvants. Frontiers in Immunology Volume 3 - 2012 (2013).

27. Case, J.B. et al. Neutralizing Antibody and Soluble ACE2 Inhibition of a Replication-Competent VSV-SARS-CoV-2 and a Clinical Isolate of SARS-CoV-2. Cell Host & Microbe 28, 475–485.e475 (2020).

28. Case, J.B. et al. Replication-Competent Vesicular Stomatitis Virus Vaccine Vector Protects against SARS-CoV-2-Mediated Pathogenesis in Mice. Cell Host Microbe 28, 465–474 e464 (2020).

29. Li, D. et al. Bacterial toxins induce non-canonical migracytosis to aggravate acute inflammation. Cell Discovery 10, 112 (2024).

30. Jiang, D. et al. Migrasomes provide regional cues for organ morphogenesis during zebrafish gastrulation. Nat Cell Biol 21, 966–977 (2019).

31. Zhang, C. et al. Monocytes deposit migrasomes to promote embryonic angiogenesis. Nat Cell Biol 24, 1726–1738 (2022).

32. Zhu, M. et al. Lateral transfer of mRNA and protein by migrasomes modifies the recipient cells. Cell Res 31, 237–240 (2021).

33. Jiao, H. et al. Mitocytosis, a migrasome-mediated mitochondrial quality-control process. Cell 184, 2896–2910 e2813 (2021).

